# AhR-mediated activation of innate lymphocytes restrains tissue-resident memory-like CD8+ T cell responses during contact hypersensitivity

**DOI:** 10.1101/2022.11.14.516493

**Authors:** S. Romero-Suárez, M.P. Correia, M. Jeong, V. Ast, M. Platten, V. Sexl, C. Mogler, A. Cerwenka, A. Stojanovic

## Abstract

Allergic contact dermatitis (ACD) and the mouse model of hapten-induced contact hypersensitivity (CHS) are inflammatory skin responses triggered by the repeated exposure to exogenous allergens and haptens. ACD and CHS effector responses have been extensively studied, but the regulatory mechanisms that control inflammation and determine the kinetics of its resolution are still incompletely understood. In addition, although CHS can be mediated by both innate and adaptive effector cells in a non-redundant manner, leading to distinct skin pathologies, their interplay during the course of inflammation remains so far unaddressed. Here, we show that NKp46^+^ innate lymphoid cells (ILCs) limit the extent of CHS inflammation by modulating the CD8^+^ T_RM_ immune compartment. This regulatory effect of ILCs depends on the expression of the ligand-induced transcription factor aryl-hydrocarbon receptor (AhR). AhR-deficiency in NKp46^+^ ILCs did not affect the memory response to hapten, but led to spatial propagation and amplification of inflammatory response in the skin. This phenotype correlated with increased numbers of *Ifng*-producing CD8^+^ T_RM_-like cells and neutrophilic infiltration in the skin. Our study thereby demonstrates a novel AhR-driven innate-adaptive immune interplay in regulating skin inflammation.

## Introduction

Allergic contact dermatitis (ACD) is a common inflammatory skin disease triggered by the repeated exposure to exogenous allergens (Vocanson et al., 2009). ACD is considered an adaptive immunity-mediated reaction, where the secondary contact with the allergen mobilizes memory T cells into the skin, and induces an inflammatory cascade involving both local and infiltrating adaptive and innate immune cells (Sheinman et al., 2021). In recent years, mounting evidence has shown that type 1 innate lymphoid cells, including natural killer (NK) cells and ILC1s, can also mediate a recall reaction in the animal model of ACD, hapten-induced contact hypersensitivity (CHS) (O’Leary et al., 2006; Sun et al., 2009; Paust et al., 2010; Peng et al., 2013; Majewska-Szczepanik et al., 2013; Wang et al., 2018; Wight et al., 2018). The recurrence and severity of ACD and CHS have been attributed to the accumulation and reactivation of resident memory CD8^+^ T (T_RM_) cells at the sites of hapten exposure (Gaide et al., 2015; Schmidt et al., 2017; Gamradt et al., 2019; Funch et al., 2022). Although effector mechanisms driving inflammation have been extensively studied, regulatory mechanisms that cease the response and determine the kinetics of its resolution are not sufficiently understood. Both innate and adaptive effector cells can mediate CHS, eliciting distinct pathological outcomes (Rouzaire et al., 2012). However, the interplay between innate and adaptive counterparts in ACD and CHS remains understudied.

Innate lymphoid cells (ILCs) play important roles in regulating skin homeostasis and inflammation (Kobayashi et al., 2020). NK cells and the three groups of ILCs (ILC1-3) comprise the innate counterparts to cytotoxic CD8^+^ T cells and helper CD4^+^ T cell system (Th1, Th2, and Th17), respectively (Spits et al., 2013). ILCs are enriched in barrier tissues and their functions are tightly regulated by cues from the tissue microenvironment, including cytokines and various metabolites (Artis and Spits, 2015). The aryl-hydrocarbon receptor (AhR) is a ligand-induced transcription factor activated by exogenous and endogenous molecules, such as dioxins, or dietary- and microbiome-derived tryptophan (Trp) metabolites (Stockinger et al., 2014). In healthy human skin and atopic dermatitis lesions, AhR is expressed by GATA3 and/or RORγt-expressing ILCs (Alkon et al., 2021), suggesting their involvement in the inflammatory response. In a murine model of CHS, ILC1s and NK cells are shown to mediate memory to haptens and reside long-term in the liver (Peng et al., 2013), where AhR ligands are abundant (Zhang et al., 2016, Fernández-Gallegos et al., 2021). Accordingly, AhR-deficient mice (AhR^−/−^) showed reduced numbers of liver-resident ILC1s, along with impaired ILC1/NK cell-mediated CHS responses (Zhang et al., 2016). However, AhR^−/−^ mice are also deficient in dendritic epidermal T cells (Kadow et al., 2011), and display a defective maturation of Langerhans cells (Jux et al., 2009), which are both implicated in the CHS reaction (Mraz et al., 2020; Jux et al., 2009). In addition to liver, “memory-bearing” ILC1s were identified in skin-draining lymph nodes 48 hours after sensitization with hapten (Wang et al., 2018). The contribution of AhR activation in the innate lymphoid system in the different phases of CHS remains enigmatic, as does the question whether AhR triggering is required to drive innate memory responses in a cell-intrinsic manner.

Here, we show that AhR activation in NKp46^+^ ILCs, comprising NK cells, ILC1s and subset of ILC3s, is dispensable for the recall response to hapten. Although ILCs could mediate memory responses independent of adaptive immunity, we demonstrate that in the presence of an intact immune system, NKp46^+^ ILCs limit the extent of CHS inflammation by modulating the CD8^+^ T_RM_-like immune compartment in an AhR-dependent manner. Thereby, we define a novel innate-adaptive immune interplay in regulating skin inflammation coordinated by innate AhR activation.

## Results and Discussion

### AhR-deficiency in NKp46^+^ ILCs results in an enhanced inflammation during CHS resolution

To investigate the role of the AhR in regulating ILC responses during CHS, we generated Ahr^f/f^Ncr1^iCre^ mice that harbor a conditional deletion of Ahr in Ncr1 (NKp46)^+^ cells, including NK cells, ILC1s and a subset of ILC3s (Eckelhart et al., 2011; Klose et al., 2014). Mice were sensitized on the back skin, and ears were challenged one week later with the hapten oxazolone (Oxa). As a measure of inflammatory response, ear thickness was assessed daily after challenge with Oxa. 24 hours post-challenge, Ahr^f/f^Ncr1^iCre^ mice presented an ear thickness comparable to the control group (Ahr^f/f^) (Figure 1A), indicating that AhR was dispensable for the generation of memory and for the development of the recall response. However, during the resolution phase, Ahr^f/f^Ncr1^iCre^ mice displayed an increased ear thickness, concomitant with the presence of confluent inflammatory sheets, increased surface of inflamed epithelium and presence of intraepithelial microabscesses in the tissue (Fig. 1B and 1C). In parallel, infiltration of mast cells and fibrosis in the inflamed ears was comparable between genotypes (Fig. S1A, B). These data indicate that, in contrast to Ahr^−/−^ mice (Zhang et al., 2016), conditional AhR deletion in the NKp46^+^ cellular compartment differentially affects the CHS response, regulating the spatial dispersion of inflammation, and that congenital AhR deficiency in other cells can indirectly affect the innate-driven memory. In addition, diverse haptens used in these studies might differentially affect innate memory through the AhR pathway. For example, it has been suggested that NK cells can directly sense haptens by the activation of Ca^2+^ permeable plasma membrane channels (Grandclément et al., 2016) and/or via MHC class I-engaging Ly49 receptors (Wight et al., 2018), which might lead to differential AhR activation.

**Figure 1.**
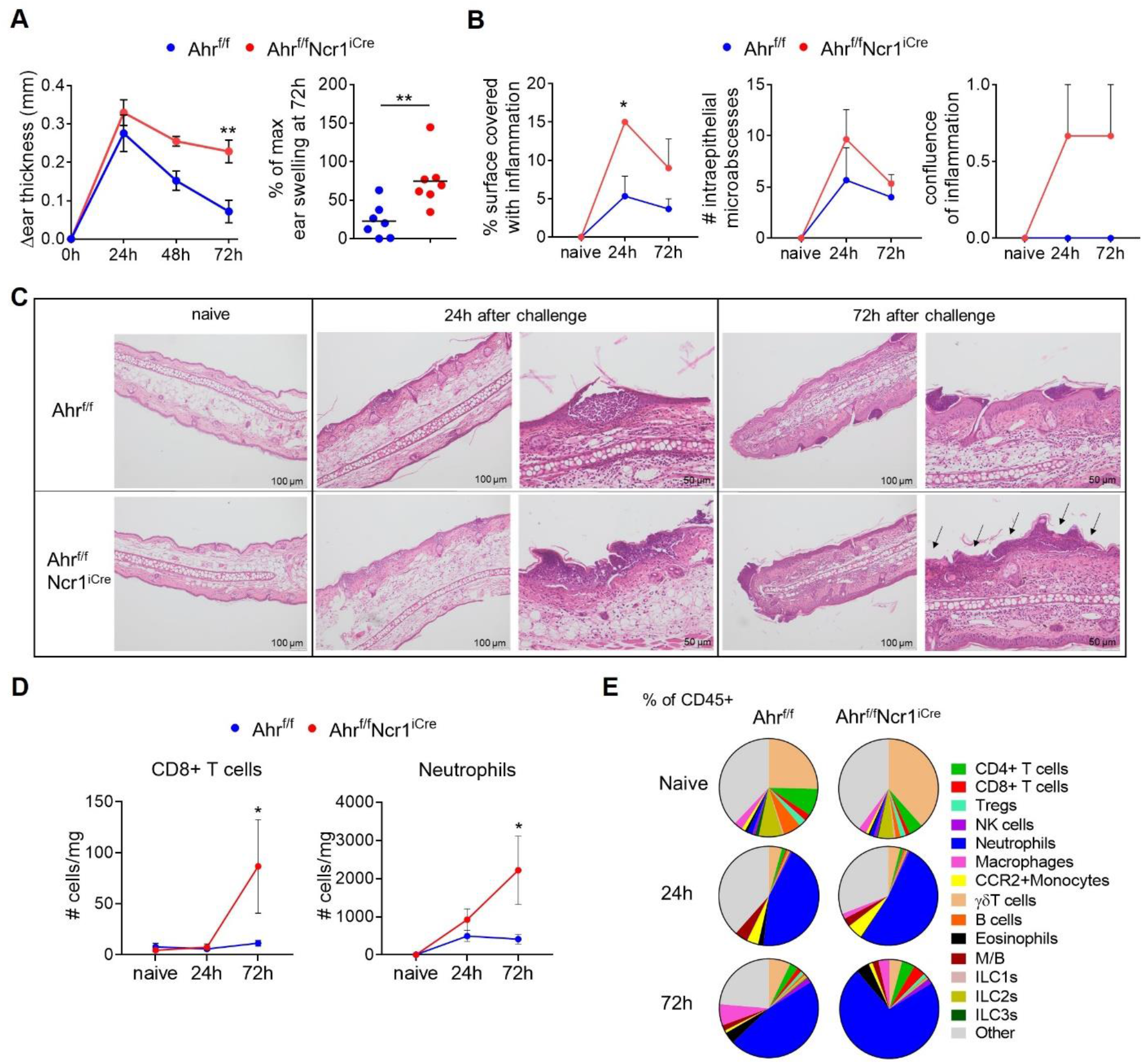
Ahr^f/f^Ncr1^iCre^ mice develop higher inflammation during CHS resolution. **A)** Change in ear thickness (relative to untreated ears) after challenge of control Ahr^f/f^ and Ahr^f/f^Ncr1^iCre^ mice with oxazolone (left, mean ± SEM, n=7, Anova with Sidak‘s post-test, **p<0.01), and % of maximum ear thickness calculated as the swelling at 72h relative to 24h after challenge (right, mean, n=7, Student t-test, **p<0.01). **B)** Pathological scoring of indicated parameters of naive/untreated and of ears 24 and 48h after challenge (n=3, mean + SEM, Anova with Sidak‘s post-test, *p<0.05). **C)** Representative H&E staining of ear sections of naive/untreated, and of ears 24 and 72h after challenge; arrows indicate confluent inflammatory infiltrate (continuous layer along epidermis). **D)** Absolute numbers (normalized to tissue weight) of CD8^+^ T cells and neutrophils in the ears of control Ahr^f/f^ and Ahr^f/f^Ncr1^iCre^ naive/untreated mice, and 24 and 72h after challenge, analyzed by flow cytometry (n=3-4, mean ± SEM, Anova with Sidak‘s post-test. *p<0.05). **E)** The mean frequencies of indicated immune cell populations among the CD45^+^ cells in the ears of control Ahr^f/f^ and Ahr^f/f^Ncr1^iCre^ naive/untreated mice, and 24 and 48h after challenge (mean, n=3-4).

To assess the quality of the CHS inflammatory response in Ahr^f/f^Ncr1^iCre^ versus control mice, we determined the composition of ear immune infiltrates (Fig. S1C-E) before and after challenge with hapten. Significantly higher numbers of CD8^+^ T cells and neutrophils were present in Ahr^f/f^Ncr1^iCre^ mice 72 hours after challenge (Fig. 1D), a time point corresponding to differential ear thickness (Fig. 1A) and an altered pathological score (Fig. 1B and 1C). Neutrophils were the dominant immune population in the inflamed ear tissue of both Ahr^f/f^Ncr1^iCre^ and control mice 24 hours after challenge (Fig. 1E), and their numbers continuously increased in Ahr^f/f^Ncr1^iCre^ mice, comprising up to 70% of all immune cells 72 hours after challenge (Fig. 1D and 1E). The numbers of other lymphoid and myeloid cell subsets did not significantly differ between Ahr^f/f^Ncr1^iCre^ and control mice (Fig. S1F). As previously reported (Paust et al., 2010), very few NK cells were present in the naive ear tissue. Upon challenge, NK cells infiltrated the inflamed skin of both Ahr^f/f^Ncr1^iCre^ and control mice, while ILC1s and ILC3s numbers remained low (Fig. S1F), indicating that AhR deficiency did not affect accumulation of these cells into inflamed ears.

Together, these data show that Ahr-deficiency in NKp46^+^ ILCs does not affect the generation of CHS memory, but spatially propagates and amplifies the inflammatory response after challenge with hapten. This phenotype correlated with increased numbers of CD8^+^ T cells and neutrophils, suggesting an AhR-driven cross-talk of innate lymphocytes to the adaptive immune system and/or to the local inflammatory mediators.

### T_RM_-like CD8^+^ T cells accumulate in the challenged skin of Ahr^f/f^Ncr1^iCre^ mice

To investigate the mechanisms that promote skin inflammation in Ahr^f/f^Ncr1^iCre^ mice, we performed single-cell transcriptome combined with feature-barcoded surface-protein expression analysis of the ear immune infiltrates 24 hours after challenge. To prevent bias towards neutrophils that dominated immune infiltrate, and to allow the detection of rare cell subsets, we enriched αβ and γδ T cell, ILC, myeloid/B-cell and the neutrophilic compartment to similar proportions in both genotypes (Fig. S2A). Unsupervised cluster analysis verified the presence of all enriched cell populations (Fig. 2A), and revealed that the CD8^+^ T cells in control mice diverged into two clusters, CD8^+^ effector T cells, which expressed *Eomes*, *Ccr7* and *Klf2*, and proliferating *Mki67*-expressing CD8^+^ T cells (Fig. 2A and S2B). In Ahr^f/f^Ncr1^iCre^ mice, we additionally identified an enrichment of CD8^+^ T cells with a core signature of skin tissue-resident T cells (Mackay et al., 2003; Pan et al., 2017; Milner et al., 2017; Kok et al., 2020), including the expression of *Gzma, Gzmb, Ifng, Klrk1* (NKG2D), *Klrc1* (NKG2A), *Ctla4, Lag3, Havcr2* (Tim-3), *Pdcd1* (PD-1), *Tnfrsf9* (CD137), *Il2ra* (CD25), *Tnfsf4* (OX40), *Cxcr6*, *Icos* and *Itgae* (CD103) (Fig. 2A, 2B and 2C). Thus, we termed this cluster “CD8^+^ T_RM_-like cells”. Comparison of the core T_RM_ signature between different CD8^+^ T cell subsets in the Ahr^f/f^Ncr1^iCre^ mice revealed overlapping signatures between proliferating and T_RM_-like CD8^+^ T cells (Fig. 2C), indicating that the accumulation of T_RM_-like CD8^+^ T cells might be the result of *in situ* differentiation/and or proliferation of the precursors in the skin.

**Figure 2.**
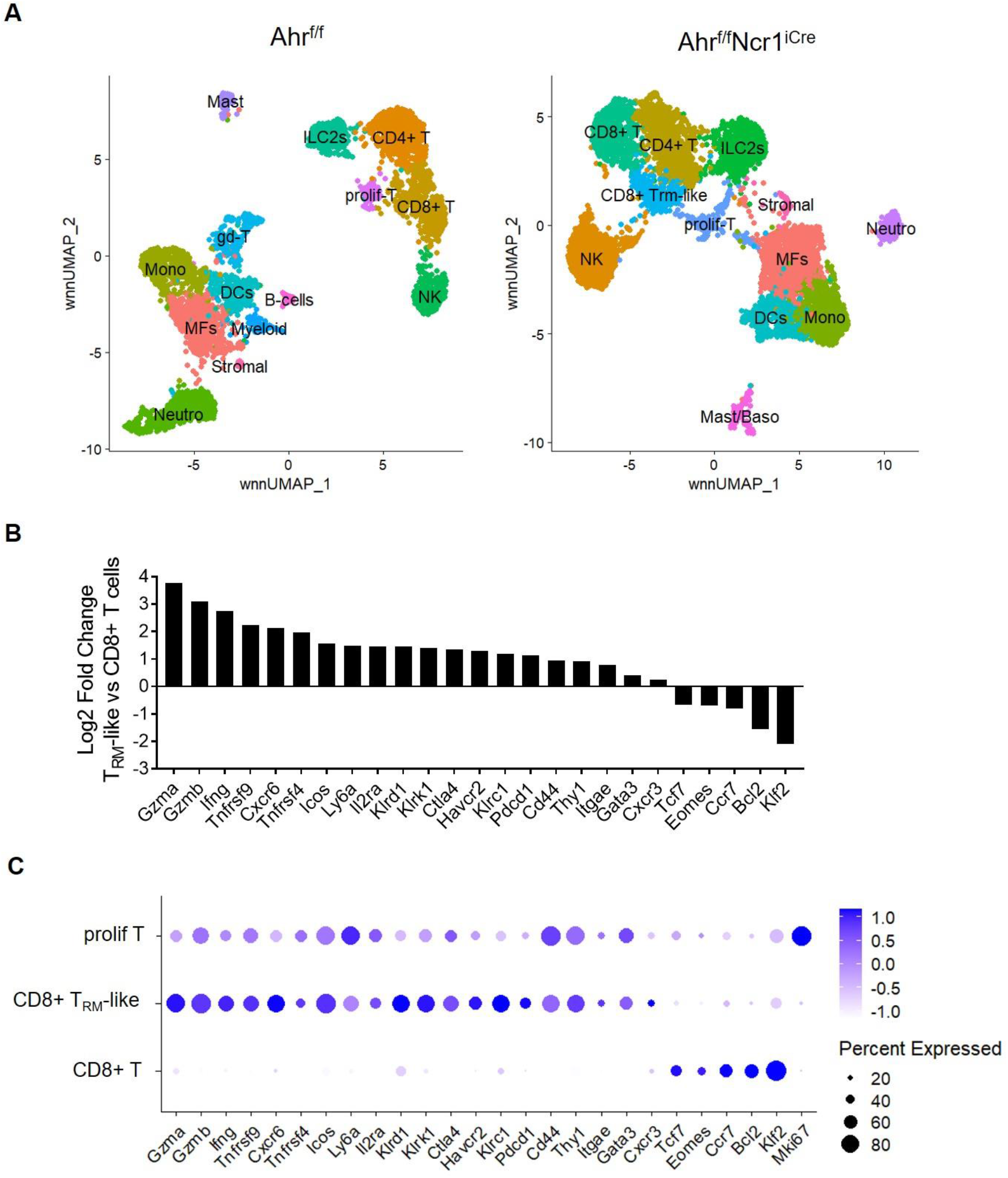
T_RM_-like CD8^+^ T cells accumulate in the challenged skin of Ahr^f/f^Ncr1^iCre^ mice. **A)** UMAP-plots showing the annotated cell clusters of single-cell sequenced immune infiltrates present in inflamed ears 24h post-challenge in Ahr^f/f^ and Ahr^f/f^Ncr1^iCre^ mice. Mast, mast cells; Mono, monocytes; Baso, basophils; Neutro, neutrophils; MFs, macrophages; prolif-T, proliferating T cells. **B)** Fold-change of the T_RM_ core-signature transcript expression between CD8^+^ T_RM_-like and CD8^+^ T cell cluster in Ahr^f/f^Ncr1^iCre^ dataset. **C)** T_RM_ core-signature transcript expression among three annotated CD8^+^ T cell subsets in the Ahr^f/f^Ncr1^iCre^ dataset.

To validate the single-cell transcriptome results, we characterized the CD8^+^ T cells present in the ear after challenge by flow cytometry. The chemokine receptor *Cxcr6* was among the top-expressed transcripts in the CD8^+^ T_RM_-like cell cluster (Fig. 2B). CXCR6 was shown to be required for T_RM_ accumulation and the acquisition of tissue-resident morphology upon treatment with the hapten DNFB (Zaid et al., 2017). We identified CXCR6-expressing CD8^+^ T cells in mice of both genotypes (Fig. 3A, left), and their proportion increased at 72 hours (resolution phase), compared to 24 hours (peak of the response) after challenge (Fig. 3A, middle) in both genotypes. However, the proportion of CXCR6^+^ CD8^+^ T cells in ears of Ahr^f/f^Ncr1^iCre^ mice was higher compared to controls, resulting in almost all CD8^+^ T cells expressing CXCR6 at 72 hours post-challenge (Fig. 3A, middle). In line with the increase of total CD8^+^ T cell numbers in the inflamed skin of Ahr^f/f^Ncr1^iCre^ mice (Fig. 1D), these data show that CD8^+^ CXCR6^+^ T cells accumulate over time in the challenged ears of Ahr^f/f^Ncr1^iCre^ mice (Fig. 3A, right). Co-expression of Ki67 (Fig. 3B, left) suggests that they are in cell cycle, and might either proliferate *in situ*, or are actively recruited upon re-activation and proliferation at distal sites, such as the lymph nodes (Gaide et al., 2015). Upon entry into the skin, T_RM_ cells progressively express CD69 and CD103 (Mackay et al., 2013), which are considered the canonical tissue-residency markers (Mueller & Mackay, 2015). In accordance with our single-cell transcriptome analysis, only a proportion of CXCR6^+^ CD8^+^ T cells co-expressed CD103 and CD69. The frequencies of these cells were similar between Ahr^f/f^Ncr1^iCre^ and control mice at 72 hours after challenge (Fig. 3B, right). The analysis of protein expression of selected transcripts from the T_RM_ core-signature (Fig. 2B), identified a unique CXCR6^+^ CD103^+^ CD8^+^ T cell subset that co-expressed Tim-3 and CD25. At 72 hours post-challenge, these cells were detected almost exclusively in Ahr^f/f^Ncr1^iCre^ mice (Fig. 3C), and might represent *bona fide* hapten-specific CD8^+^ T_RM_ cells.

**Figure 3.**
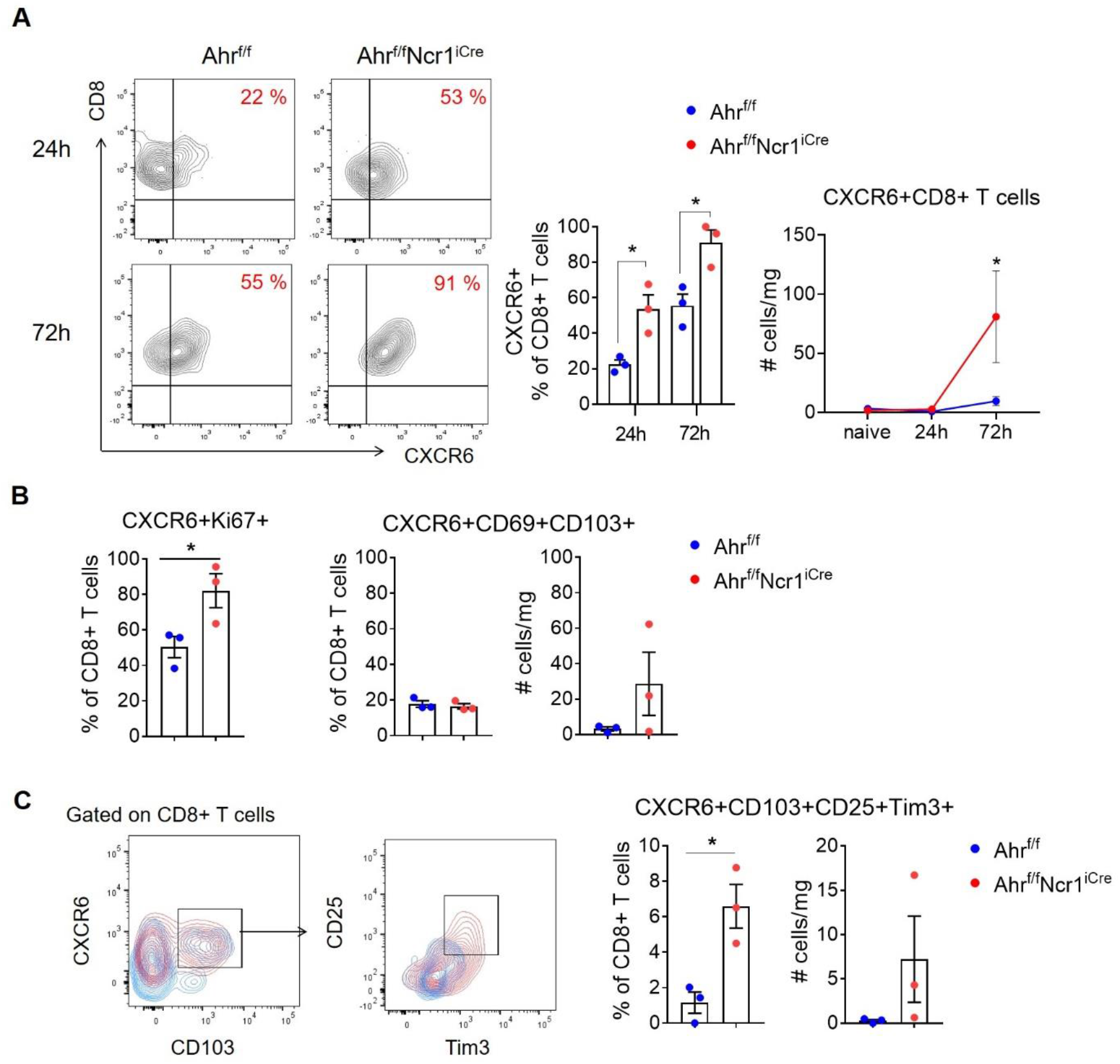
CXCR6-expressing T_RM_-like CD8^+^ T cells accumulate in the inflamed ear skin of Ahr^f/f^Ncr1^iCre^ mice. **A)** Representative contour-plots (left) showing the CXCR6 expression on gated CD8^+^ T cells in the ears of Ahr^f/f^ and Ahr^f/f^Ncr1^iCre^ mice 24h and 72h after challenge. Percentage of CXCR6^+^ cells among CD8^+^ T cells (middle, n=3, mean + SEM, Anova with Sidak’s post-test *p<0.05) and the absolute numbers of CXCR6^+^ CD8^+^ T cells normalized to tissue weight (right, n=3, mean ± SEM, Anova with Sidak’s post-test, *p<0.05). **B)** Percentage of CD8^+^ CXCR6^+^ T cells expressing Ki67 (left, n=3, mean ± SEM, Student’s t-test, *p<0.05) or tissue-residency markers CD103 and CD69 (middle) 72h post-challenge. Absolute cell numbers of CXCR6^+^ CD103^+^ CD69^+^ CD8^+^ T cells normalized to tissue weight (right, n=3, mean ± SEM, Student’s t-test, *p<0.05). **C)** Representative contour-plots showing CXCR6, CD103, CD25 and Tim-3 expression on gated CD8^+^ T cells from the inflamed ears of Ahr^f/f^ (blue) and Ahr^f/f^Ncr1^iCre^ (red) mice 72h post-challenge (left). Frequency and absolute cell numbers of CXCR6^+^ CD103^+^ CD25^+^ Tim-3^+^ CD8^+^ T cells normalized to tissue weight (right, n=3, mean ± SEM, Student’s t-test, *p<0.05)

### CD8^+^ T_RM_-like cells express Ifng in the inflamed ears of Ahr^f/f^Ncr1^iCre^ mice

To understand how T_RM_-like CD8^+^ T cells contribute to the increased inflammation in Oxa-challenged Ahr^f/f^Ncr1^iCre^ mice, we first analyzed their cytokine-expression profile. CHS studies using different haptens have reported that CD8^+^ T_RM_ cells promote CHS inflammation via IFN-γ and IL-17 secretion (Kish et al., 2009; He et al., 2009; Kawano et al., 2014; Chong et al., 2014). T_RM_-like CD8^+^ T cells in ears challenged with Oxa expressed predominantly *Ifng*, with some cells also transcribing *Csf1* and/or *Areg* (Fig. 4A), encoding IFN-γ, M-CSF and amphiregulin, respectively. Some of the remaining CD8^+^ T cells in both control and Ahr^f/f^Ncr1^iCre^ mice also expressed these transcripts (Fig. 4A). As these cytokines, can also be produced by other immune cells (Monticelli et al., 2015; Burzyn et al., 2013; Ushach & Zlotnik, 2016), we assessed the relative contribution of different cell subsets to the overall expression of these cytokines, as well as their relative enrichment in the whole immune compartment in control and Ahr^f/f^Ncr1^iCre^ mice. Out of the three selected candidates, *Ifng* was the only transcript enriched in the immune infiltrate of Ahr^f/f^Ncr1^iCre^ mice (Fig. 4B). *Csf1* was mainly transcribed by myeloid and stromal cells, while γδ T cells, ILC2s and some cells within the proliferating CD8^+^ T cluster expressed *Areg* (Fig. S2C). In control animals, NK cells were the main producers of *Ifng*, while in Ahr^f/f^Ncr1^iCre^ mice, T_RM_-like and proliferating T cells also contributed to *Ifng* expression (Fig. 4C). It was recently shown that CD8^+^ T_RM_ cells mediated neutrophil recruitment by inducing CXCL1 and CXCL2 production in the skin (Funch et al., 2022). We found that *Ifngr2*, encoding the IFN-γ receptor 2, was preferentially expressed by the stromal and myeloid compartment, which also expressed the transcripts *Cxcl1*, *Cxcl2* and *Cxcl3*, reported as neutrophil-attracting chemokines (Dilulio et al., 1999; Engeman et al., 2004; Kish et al., 2012) (Fig. 4D). Overall, these data suggest that the T_RM_-like CD8^+^ T cells that accumulate in the inflamed ears of Ahr^f/f^Ncr1^iCre^ mice significantly contribute to the increased availability of IFN-γ in the tissue. IFN-γ-sensing and neutrophil-attracting chemokine production seems to be integrated by the myeloid/stromal compartment, resulting in an increased neutrophilic infiltration during CHS.

**Figure 4.**
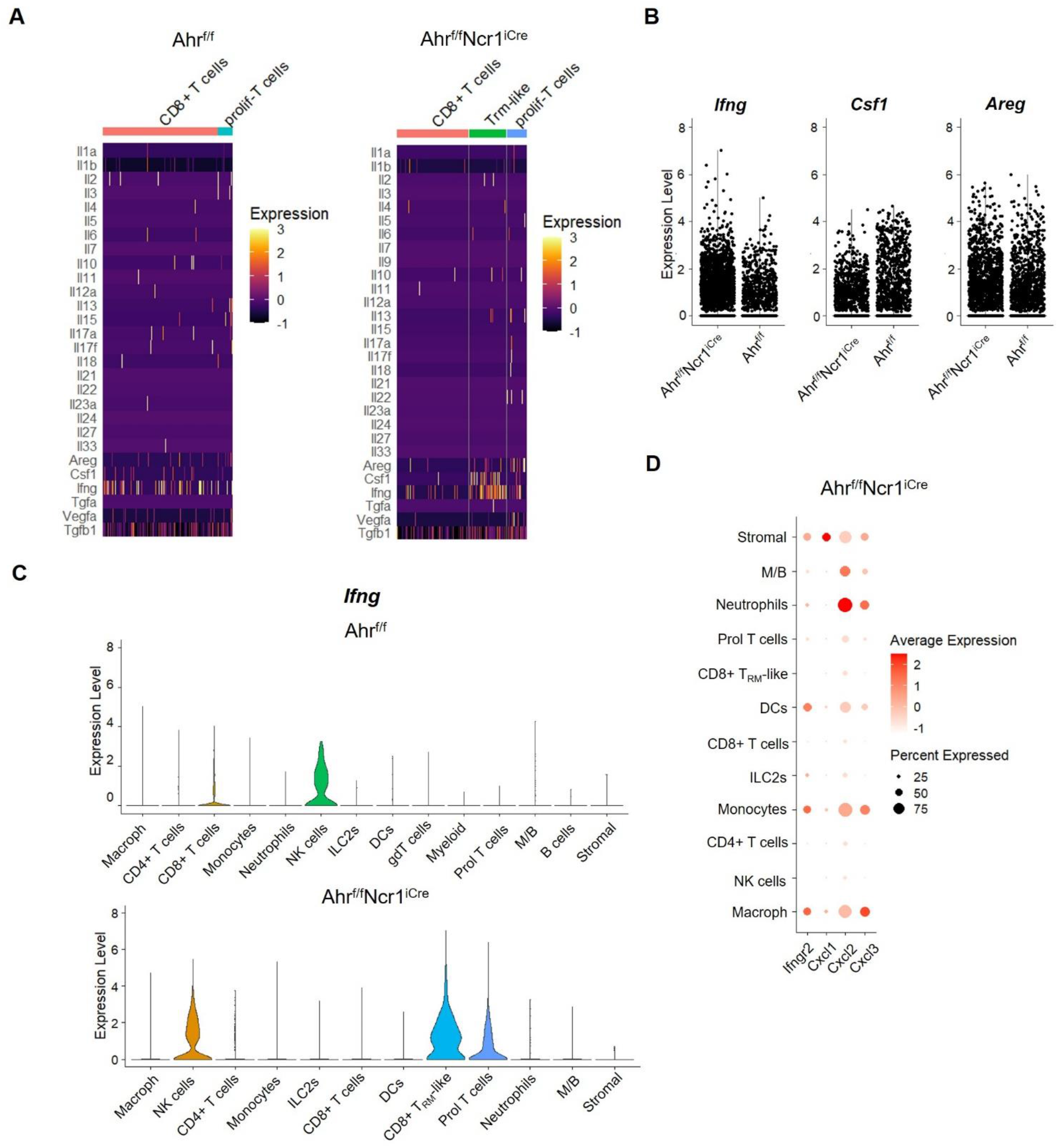
CD8^+^ T_RM_-like cells express *Ifng* in the inflamed ears of Ahr^f/f^Ncr1^iCre^ mice. **A)** Heatmap of normalized cytokine transcript expression in single-cells of the indicated CD8^+^ T cell clusters in Ahr^f/f^ and Ahr^f/f^Ncr1^iCre^ mice 24h post-challenge. **B)** Expression of *Ifng*, *Csf1* and *Areg* in the immune cell compartment of Ahr^f/f^Ncr1^iCre^ and Ahr^f/f^ mice, each dot represents a cell. **C)** Violin plots of *Ifng* expression along annotated cell clusters in Ahr^f/f^ and Ahr^f/f^Ncr1^iCre^ mice 24h post-challenge. **D)** Expression of *Ifngr2* and neutrophil-attracting chemokine transcripts along annotated cell clusters in Ahr^f/f^Ncr1^iCre^ mice 24h post-challenge. M/B, mast cells and basophils

To determine whether the adaptive immune compartment transmits the signal downstream of AhR-deficient innate lymphocyte responses to immediate inflammatory mediators in the skin, we generated Rag2-deficient Ahr^f/f^Ncr1^iCre^ mice. In accordance with previously published data (O’Leary et al., 2006; Paust et al., 2010; Majewska-Szczepanik et al., 2013; Peng et al., 2013; Wang et al., 2018), an innate memory response to Oxa was generated in the absence of adaptive immune cells (Fig. 5). Both, Rag2^−/−^Ahr^f/f^Ncr1^iCre^ mice and control Rag2^−/−^Ahr^f/f^ mice mounted a similar CHS response 24 hours after challenge. In contrast to Ahr^f/f^Ncr1^iCre^ mice, Rag2-deficient Ahr^f/f^Ncr1^iCre^ mice did not present an increased ear thickness during the resolution phase. These data show that AhR-deficient innate lymphocytes, although able to mediate a memory response, are not sufficient to drive an increased inflammation in the absence of adaptive immune cells.

**Figure 5.**
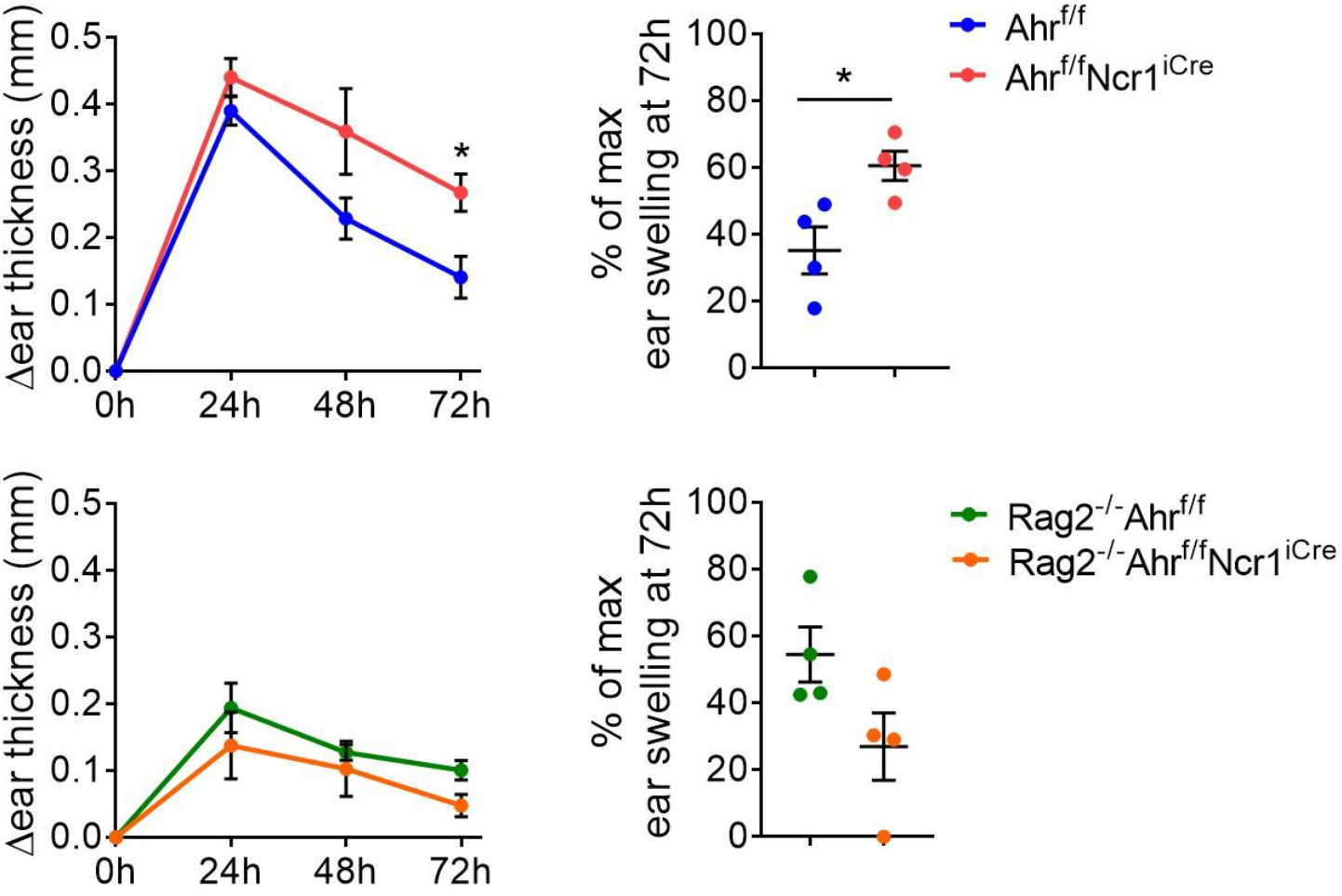
Adaptive immune cells mediate increased CHS inflammation in Ahr^f/f^Ncr1^iCre^ mice. CHS response in Ahr^f/f^ and Ahr^f/f^Ncr1^iCre^ mice (upper panel), or Rag2^−/−^Ahr^f/f^ and Rag2^−/−^ Ahr^f/f^Ncr1^iCre^ mice (lower panel), depicted as the change in ear thickness (relative to untreated ears) after challenge with oxazolone (left), and as the percentage of remaining ear swelling at 72h relative to the maximum of response (24h post-challenge) (right). Ear thickness curves were analyzed using an Anova with Sidak‘s post-test, and the maximum of ear swelling with a Student’s t-test; *p<0.05. n=4 mice per group, mean ± SEM.

### Repeated challenge with hapten leads to the enrichment of CD8^+^ T_RM_-like cells and increased inflammatory score of CHS

CD8^+^ T_RM_ cells are shown to accumulate at the sites of previous hapten exposures correlating with the magnitude of the subsequent allergic reaction (Gaide et al., 2015; Gamradt et al., 2019; Funch et al., 2022). To test if AhR-deficiency in NKp46-expressing cells affects the magnitude of T_RM_ cell accumulation in these settings, we exposed sensitized Ahr^f/f^Ncr1^iCre^ mice to three consequent weekly challenges with Oxa (Fig 6A, left). Both control and Ahr^f/f^Ncr1^iCre^ mice mounted similar maximal responses, measured by ear thickness (Fig. 6A, right). Upon 3^rd^ challenge, CD8^+^ T_RM_-like cells were detected in the inflamed ears of mice in both genotypes; however, their numbers were significantly higher in Ahr^f/f^Ncr1^iCre^ mice (Fig. 6B). Despite presenting comparable ear thickness to controls animals, Ahr^f/f^Ncr1^iCre^ mice displayed higher neutrophilic infiltrates (Fig. 6B) and more severe pathological score, including higher surface covered with inflammatory cells and higher confluence of inflammation (Fig. 6C). Therefore, AhR-deficiency in the innate lymphoid compartment, which results in an increased inflammation initially observed in the resolution phase after primary challenge, leads to the incremental amplification of T_RM_-driven responses upon repeated challenge, worsening skin pathology. In sum, the innate lymphocyte-AhR system acts as a constraint-point that limits the allergic skin inflammatory reaction.

**Figure 6.**
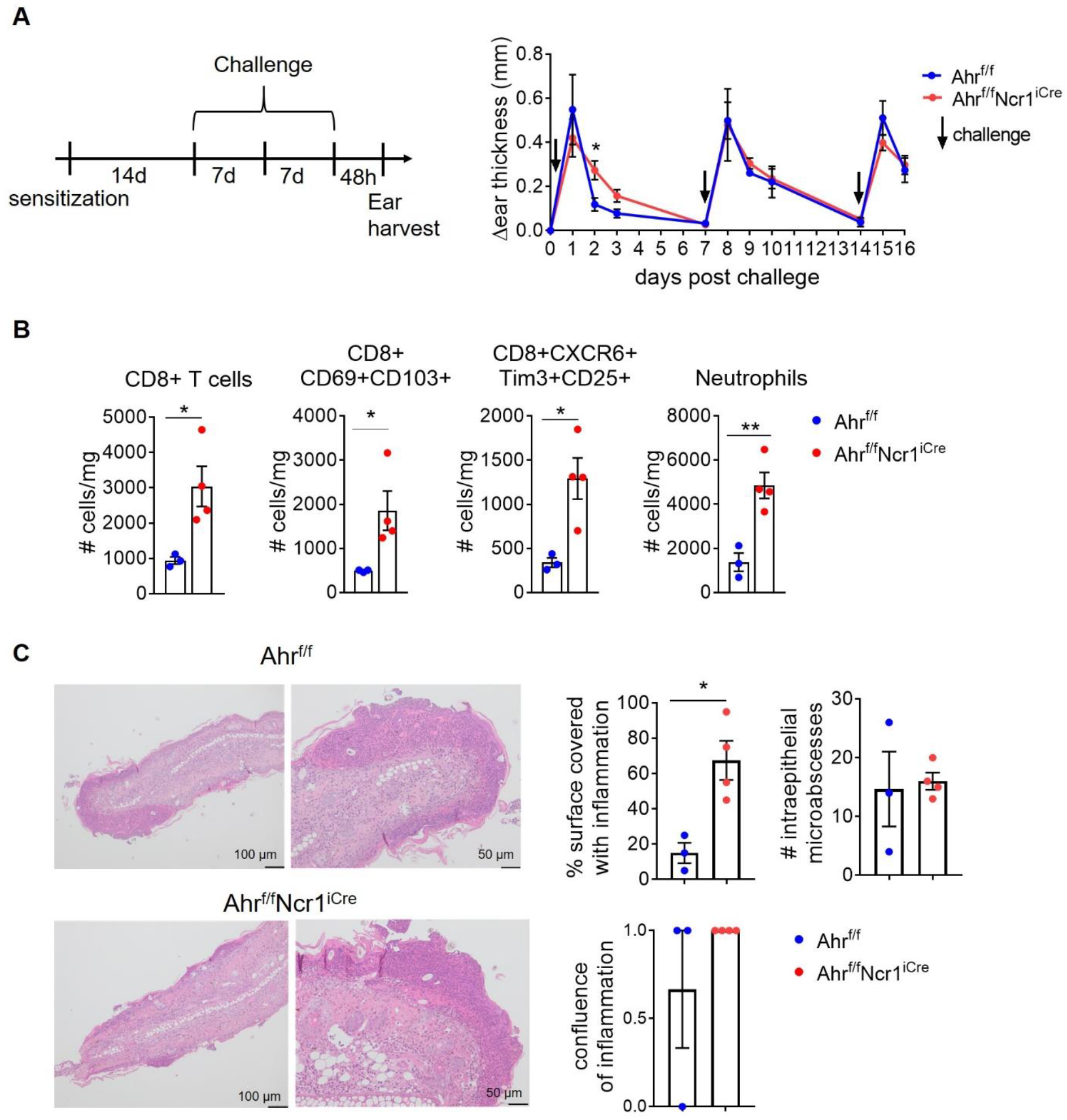
Repeated challenge with hapten leads to an enrichment of CD8^+^ T_RM_-like cells and increased inflammatory score in Ahr^f/f^Ncr1^iCre^ mice. **A)** Experimental setup of repeated challenge (left) and the corresponding analysis of response depicted as the change in ear thickness (relative to untreated ears) after challenge with oxazolone (right). **B)** Absolute numbers (normalized to tissue weight) of total CD8^+^ T cells, CD103^+^ CD69^+^ CD8^+^ T cells, CXCR6^+^ Tim-3^+^ CD25^+^ CD8^+^ T cells, and neutrophils in the ear 48h after the third challenge (n=3-4, mean ± SEM, Student’s t-test, *p<0.05, **p<0.01). **C)** Representative H&E staining of ear tissue harvested 48h after the third challenge (left), and histopathological scoring (right); n=3-4 mean ± SEM, Student’s t-test, *p<0.05.

## Conclusions

In immunocompetent organisms, the innate and adaptive immune system operate together to mount efficient, but controlled immune responses and enable protection against pathogens or transformed cells. Early activation of innate immunity is considered indispensable to provide microenvironmental contexture and to instruct the type of adaptive immune response needed against different types of pathogens in different microenvironments. For example, secretion of IFN-γ by NK cells in the lymph node was reported essential for priming Th1 immunity by dendritic cells (Martin-Fontecha et al., 2004). Equally important to launching and propelling the adequate response, is the ability to cease it and control it. So far, the upmost relevance to these functions was attributed to regulatory cells with the ability to dampen the inflammatory response and mediate tissue repair, such as regulatory T cells or M2-like macrophages, respectively. Here, we identified a so far unappreciated role of innate lymphocytes to control the inflammatory response in the skin evoked by (repeated) challenge with hapten. Inability of NKp46^+^ ILCs to engage AhR activation led to the amplification of T_RM_-driven inflammatory responses in the hapten-challenged skin. The increased numbers of T_RM_-like cells correlated with augmented neutrophilic infiltration and resulted in the formation of continuous inflammatory sheets along the epidermis, thus worsening skin pathology. It remains to be investigated which subset of NKp46-expressing ILCs mediates these effects, and if congenital deletion of AhR inherently changes the nature of their cross-talk with the adaptive immune system, or alternatively, if the inflammation-induced AhR activation is required to control the quality and/or quantity of CD8^+^ T cell activation in context of allergic dermatitis. AhR activation and AhR ligands play important roles in regulating skin homeostasis and immune responses (Fernández-Gallegos et al., 2021). Although it is plausible that local exposure to AhR ligands might regulate the ILC-T cell cross-talk in the skin, memory ILCs were found to reside in the liver (Peng et al., 2013), where the concentration of AhR ligands, such as Kynurenine, is high (Bunger et al., 2008). Therefore, it is attractive to speculate that their functional competence to regulate T cell responses might also be shaped and/or maintained in the liver. In addition, congenital absence of NKp46-expressing ILC3s that depend on AhR signaling for their development in the gut (Kiss et al., 2011), also suggests a conceivable role of a gut-skin axis in the regulation of skin immune responses. The AhR-ILC module may be also required to bridge the local skin-resident commensals to T cell function during inflammation, in line with findings demonstrating the requirement of skin microbiome for optimal skin immune fitness, through modulation of T cell effector output (Naik et al., 2012).

Augmentation of T_RM_ differentiation, recruitment or proliferation regulated by the AhR-ILC axis has multiple implications for skin disease. Regulating T cell responses could be of high relevance not only for allergic, but also for other types of dermatitis, which presents the highest global disease burden among skin diseases, and significantly affects the quality of life of affected individuals (Sheinman et al., 2021). T_RM_ cells in particular, were reported to promote skin inflammation, rapid development of flares, relapse, and chronic responses, including vitiligo and psoriasis (Vo et al., 2019; Gallais Sérézal et al., 2019; Richmond et al., 2019; Boniface et al., 2018). As they exert both residency and memory, and their numbers incrementally increase with every subsequent exposure to allergen, they bear a colossal pathological potential. Cell-intrinsic mechanisms based on expression of immunoregulatory receptors, such as PD-1 or Tim-3, have been proposed to curtail T_RM_ re-activation, and could be exploited as therapeutic targets (Gamradt et al., 2019). However, insight into cell-extrinsic regulatory mechanisms able to skew T cells responses away from T_RM_ establishment in the skin are desirable to further improve management of patients with dermatitis. Here, we propose that the AhR-ILC system is able to fine-tune T cell responses in the skin, and therefore, might represent a unique early check-point to regulate the outcome of skin inflammation.

## Materials and Methods

### Mice

Mice were bred and housed in the animal facility of the Medical Faculty Mannheim, University of Heidelberg, under specific pathogen-free conditions. C57BL6/N mice were purchased from Janvier. AhR^f/f^ mice (Ahr^tm3.1Bra^; Walisser et al., 2005) were crossed with Ncr1^iCreTg^ mice (Tg-Ncr1-iCre^265Sxl^; Eckelhart et al., 2011) to generate AhR^f/f^Ncr1^iCreTg^ mice. Rag2^−/−^ mice (B6.129S-Rag2^tm1Fwa^), bred in house, were crossed to AhR^f/f^Ncr1^iCreTg^ mice to generate Rag2^−/−^AhR^f/f^Ncr1^iCreTg^ mice. Both female and male animals of an age 10 to 32 weeks were used in the experiments. All animal experiments were approved by the local animal welfare commission at the Regierungspräsidium Karlsruhe.

### Hapten-induced contact hypersensitivity

Mice were sensitized on the shaved back-skin for two consecutive days with 50 μl of 5% Oxa (4-Ethoxymethylen-2-phenyl-2-oxazolin-5-on, Sigma-Aldrich) diluted in methanol and acetone (1:1, v/v). One to two weeks after sensitization, mice were challenged by the application of 2% Oxa in the dorsal and frontal side of the ear (10 μl each side). Ear thickness was measured with a caliper before challenge, and every 24 hours after challenge. For repeated challenge, ears were treated every 7 days for three consecutive weeks.

### Histological analysis

Ear tissue was fixed in 4% paraformaldehyde at 4°C overnight. Tissue was then rinsed with tap water, dehydrated in ethanol series (75-100%) and xylene, and embedded in paraffin. Paraffin sections (2μm) were cut and stained with hematoxylin and eosin (H&E), toluidine-blue or Masson’s trichrome staining, according to standard protocols. The histopathological evaluation was performed in a blinded fashion. Percentage of surface covered by inflammation was estimated by semiquantitative analysis. Confluence of inflammation was scored as 1 when a continuous infiltration of the epidermis by granulocytes was present, or as 0 when absent (Fig. S3). The number of intraepithelial microabscesses was counted in 5 high-power fields and over the whole ear surface. Pictures were taken with an Olympus BX 46 microscope and the Olympus UC90 camera (software: cellSens Entry, Olympus).

### Preparation of single-cell suspensions from ear tissue

The ventral and dorsal part of the ears were separated, minced and digested using the Tissue Dissociation Kit 1 and a gentleMACS™ Dissociator (both from Miltenyi Biotec). Cells were filtered through a 70 μm cell strainer, and washed twice in PBS/10% FCS (ThermoFisher) before staining.

### Flow cytometry analysis and cell sorting

Cells were stained with Zombie Aqua™ Fixable Viability Kit (Biolegend), to exclude apoptotic and death cells. Fc receptors were blocked with anti-mouse CD16/32 (TruStain FcX™, Biolegend). Cells were then incubated with fluorochrome-conjugated antibodies for 20 min at 4°C and washed twice with PBS. Intracellular staining was performed using the eBioscience™ Foxp3/Transcription Factor Staining Buffer Set (Invitrogen). Cells were acquired with a LSRFortessa™, and data were analyzed using FlowJo™ software (both from BD Biosciences). For single-cell transcriptome/surface protein analysis, cell suspension from hapten-challenged (24 hours) ear tissue were prepared from 10 individual mice per genotype. Cells were pooled, and then incubated with a mixture of fluorophore-conjugated and TotalSeq™-B oligo-conjugated antibodies (Biolegend). After washing, immune cells were enriched using flow-cytometric sort (BD FACSAria™ Fusion) yielding relative frequencies of approximately 20% T, 30% NK/ILCs, 10% DETCs/γδT, 10% neutrophils and 30% myeloid/B cells, to allow detection of subpopulations and/or rare cells by multiomic data analysis. Cells were gated as single/live CD45-expressing T cells (Ly6G^neg^TCRγδ^neg^TCRβ^+^), NK cells and ILCs (TCRβ^neg^TCRγδ^neg^CD90.2^+/neg^NK1.1^+/neg^), DETCs and γδ T cells (Ly6G^neg^TCRβ^neg^TCRγδ^+^), neutrophils (Ly6G^+^), and myeloid and B cells (Ly6G^neg^TCRβ^neg^TCRγδ^neg^CD90.2^neg^NK1.1^neg^) (Fig S2A). Dead cells were excluded using 7-AAD (Biolegend).

### Single-cell RNA and feature-barcoded surface protein library preparation

Single-cell Gel-bead-in-Emulsions (GEMs) and mRNA libraries were generated by the Next-Generation Sequencing (NGS) Core Facility of UMM, using 10x Genomics Chromium single-cell 3’ library and gel bead kit v3.1 and Dual Index Kit Set A, according to manufacturer’s instructions. A total of 10000 cells per sample/genotype were loaded into a Chromium Next GEM Chip G (10x Genomics). 3’ Gene Expression Library and Cell Surface Protein Library were mixed in 4:1 ratio, and sequenced on Illumina NextSeq 500/550 using High Output Kit v2.5 (150 Cycles).

### Single-cell multiomic data analysis

Single-cell datasets were analyzed and visualized using the Seurat R package (version 4.0.3) (Stuart et al., 2019). For quality control, transcripts, which were detected in less than three cells, and cells with more than 6000 and less than 200 genes with non-zero counts, were filtered out. All cells having more than 20% of mitochondrial gene counts were excluded. For each dataset, gene expression and protein data were integrated and the clustering of cells was performed using the weighted-nearest neighbor method (Hao et al., 2021). Cell clusters were annotated based on differential transcript and protein expression as: macrophages (*Mrc1, Mafb, Ctsl* and F4/80); CD4^+^ T cells (*Cd4, Trbc2, Cd3d*, CD3 and CD4); CD8^+^ T cells (*Cd8b1 Cd8a Cd3d*, CD3 and CD8); monocytes (*Ly6c2, Lyz2, Plac8* and Ly6C); neutrophils (*S100a9*, *S100a8, Hcar2*); NK cells (*Eomes, Gzma, Prf1, Klra8, Klra4, Klre1* and NKp46); ILCs (*Rora, Ikzf3, Areg, Gata3, Ly6a* and CD127); dendritic cells (DCs) (*H2-DMb1, H2-Eb1, H2-Aa, H2-Ab1, H2-DMa, Cd74*, MHC-II and CD11c); DETCs/γδT cells (*Tcrg-C1, Trdc, Cd3e, Cd3g* and CD3); myeloid cells (*Cd74, Tmem123, Ccr7* and MHC-II); mast cells and basophils (M/B) (*Gata2, Cpa3 Mcpt4, Mcpt8* and FcεRIα); proliferating T cells (*Mki67*, CD3, CD4 and CD8); B cells (*Ebf1, Ms4a1 and* MHC-II); and stromal cells (*Col8a1, Igfbp7, Col1a2, Sparc, Mgp*).

### Statistical analysis

Results are reported as mean ± SEM, n represents numbers of animals per experimental group. Using GraphPad Prism, data were tested for normal distribution with Shapiro-Wilk test, followed by statistical test corrected for multiple comparisons when necessary. Experimental groups were considered significantly different when *, p<0.05, **, p<0.01, ***, p<0.001.

### Antibodies used for flow cytometry, cell sorting and CITE-seq

| **Antibody** | **Clone** | **Company** |
| --- | --- | --- |
| anti-mouse CCR2 BUV496 | 475301 | BD Biosciences |
| anti-mouse CD11b BV650 | M1/70 | Biolegend |
| anti-mouse CD11c PerCP/Cy5.5 | N418 | Biolegend |
| anti-mouse CD19 PE/Cy7, APC/Cy7 | 6D5 | Biolegend |
| anti-mouse CD19 BUV563 | 1D3 | BD Biosciences |
| anti-mouse CD127 APC/Cy7 | A7R34 | Biolegend |
| anti-mouse CD103 BUV395 | 2E7 | BD Biosciences |
| anti-mouse CD137 APC | 17B5 | Biolegend |
| anti-mouse CD25 PE/Dazzle 594, BV650 | PC61 | Biolegend |
| anti-mouse CD3 PE/Cyanine7, APC/Fire 750 | 17A2 | Biolegend |
| anti-mouse CD3e BUV496 | 500A2 | BD Biosciences |
| anti-mouse CD4 BUV563 | GK1.5 | BD Biosciences |
| anti-mouse CD8a FITC, BV421 | 53-6.7 | Biolegend |
| anti-mouse CD45 BV785 | 30-F11 | Biolegend |
| anti-mouse CD69 PerCP/Cy5.5 | H1.2F3 | Biolegend |
| anti-mouse CD90.2 FITC | 53-2.1 | Biolegend |
| anti-mouse CXCR6 Alexa Fluor 647, BV421 | SA051D1 | Biolegend |
| anti-mouse Eomes PE-CF594 | X4-83 | BD Biosciences |
| anti-mouse FcεR1α APC/Cy7 | MAR-1 | Biolegend |
| anti-mouse Foxp3 PE-CF594 | MF23 | BD Biosciences |
| anti-mouse F4/80 PE-CF594 | T45-2342 | BD Biosciences |
| anti-mouse F4/80 APC/Cy7 | BM8 | Biolegend |
| anti-mouse GATA3 Alexa Fluor 647 | 16E10A23 | Biolegend |
| anti-mouse I-A/I-E APC, APC/Cy7 | M5/114.15.2 | Biolegend |
| anti-mouse Ki67 PE/Cy7 | 16A8 | Biolegend |
| anti-mouse Lag3 PE | C9B7W | Biolegend |
| anti-mouse Ly6C FITC | HK1.4 | Biolegend |
| anti-mouse Ly6G BUV805 | 1A8 | BD Biosciences |
| anti-mouse Ly6G APC/Cy7 BV421 | 1A8 | Biolegend |
| anti-mouse NKp46 PE/Cy7, APC/Cy7 | 29A1.4 | Biolegend |
| anti-mouse NK1.1 BV650, BV711 | PK136 | Biolegend |
| anti-mouse PD1 BUV737 | J43 | BD Biosciences |
| anti-mouse RORγt PE | Q31-378 | BD Biosciences |
| anti-mouse Siglec-F PE | E50-2440 | BD Biosciences |
| anti-mouse T-bet PerCP/Cy5.5 | 4B10 | BD Biosciences |
| anti-mouse TCRβ BUV805 | H57-597 | BD Biosciences |
| anti-mouse TCRβ BV711 | H57-597 | Biolegend |
| anti-mouse TCR$\gamma\delta$ BV711 | GL3 | BD Biosciences |
| TotalSeq™-B0182 anti-mouse CD3 | 17A2 | Biolegend |
| TotalSeq™-B0001 anti-mouse CD4 | RM4-5 | Biolegend |
| TotalSeq™-B0002 anti-mouse CD8a | 53-6.7 | Biolegend |
| TotalSeq™-B0184 anti-mouse CD335 | 29A1.4 | Biolegend |
| TotalSeq™-B0198 anti-mouse CD127 | A7R34 | Biolegend |
| TotalSeq™-B0013 anti-mouse Ly6C | HK1.4 | Biolegend |
| TotalSeq™-B0115 anti-mouse FcεR1α | MAR-1 | Biolegend |
| TotalSeq™-B0106 anti-mouse CD11c | N418 | Biolegend |
| TotalSeq™-B0117 anti-mouse I-A/I-E | M5/114.15.2 | Biolegend |
| TotalSeq™-B0114 anti-mouse F4/80 | BM8 | Biolegend |

## Supporting information

RomeroSuarez-et-al_Supplement

## Acknowledgements

We thank Angelika Duda and Tina Fuchs for assistance with multiomics experiments; Petra Bugert, Marian Wincher and Patrick Matei for technical support; Sophia Papaioannou for assistance with experiments; the animal facility of the Medical Faculty Mannheim and the German Cancer Research Center for assistance with animal care and experiments; ZMF Mannheim for supporting the processing of biological samples; and the members of SFB TRR 156 for helpful discussion.

## Funding

The project was supported by grants from the German Research Foundation [SFB-TRR156 (B10N to AC), SFB1366 (Project number 319 394046768-SFB 1366; C02 to AC, C01 to MP and Z2 to CM), SPP1937 (CE 140/2-2 to AS and AC), RTG2727 – 445549683 Innate Immune checkpoints in cancer and tissue damage (B1.2 to AC and AS; B1.1 to MP;)]; by a network grant of the European Commission (H2020-MSCA-MC322, ITN-765104-MATURE-NK); and by the Angioformatics platform of the European Center for Angioscience (ECAS).

## Author Contribution

SRS performed experiments, analyzed the data and wrote the manuscript; MPC performed experiments, analyzed the data, and provided critical input and ideas; MJ performed experiments; VA analyzed single-cell multiomics data; MP provided animals and critical input; VS provided animals, CM performed histological analyzes; AC supervised the project and wrote the manuscript; AS conceptualized and supervised the project, performed experiments, analyzed the data and wrote the manuscript.

## Declaration of interests

The authors declare no competing interests.

## Notes

### Competing Interest Statement

The authors have declared no competing interest.

