## Supplementary material for "AhR-mediated activation of innate lymphocytes restrains tissue-resident memory-like CD8+ T cell responses during contact hypersensitivity": RomeroSuarez-et-al_Supplement

[1]Department of Immunobiochemistry, Mannheim Institute for Innate Immunoscience (MI3), Medical Faculty Mannheim, Heidelberg University, Mannheim, Germany
[2]Cancer Biology and Epigenetics Group, Research Center of IPO Porto (CI-IPOP)/RISE@CI-IPOP (Health Research Network), Portuguese Oncology Institute of Porto (IPO Porto)/Porto Comprehensive Cancer Center (Porto.CCC), Porto, Portugal
[3]NGS Core Facility, Medical Faculty Mannheim, Heidelberg University, Mannheim, Germany
[4]Institute for Clinical Chemistry, University Hospital Mannheim (UMM), Medical Faculty Mannheim, Heidelberg University, Mannheim, Germany
[5]CCU Neuroimmunology and Brain Tumor Immunology, German Cancer Research Center (DKFZ), German Cancer Consortium (DKTK), Clinical Cooperation Unit (CCU), Heidelberg, Germany
[6]Department of Neurology, University Hospital Mannheim (UMM), MCTN, Medical Faculty Mannheim, Heidelberg University, Mannheim, Germany
[7]DKFZ Hector Cancer Institute at the University Medical Center Mannheim, Mannheim, Germany
[8]Department for Biomedical Sciences, Institute of Pharmacology and Toxicology, University of Veterinary Medicine, Vienna, Austria
[9]Institute of Pathology, School of Medicine, Technical University Munich, Munich, Germany
[10]European Center for Angioscience (ECAS), Medical Faculty Mannheim, Heidelberg University, Mannheim, Germany
[#]Equally contributing senior authors

### *Supplementary Figures*

### *Figure S1*

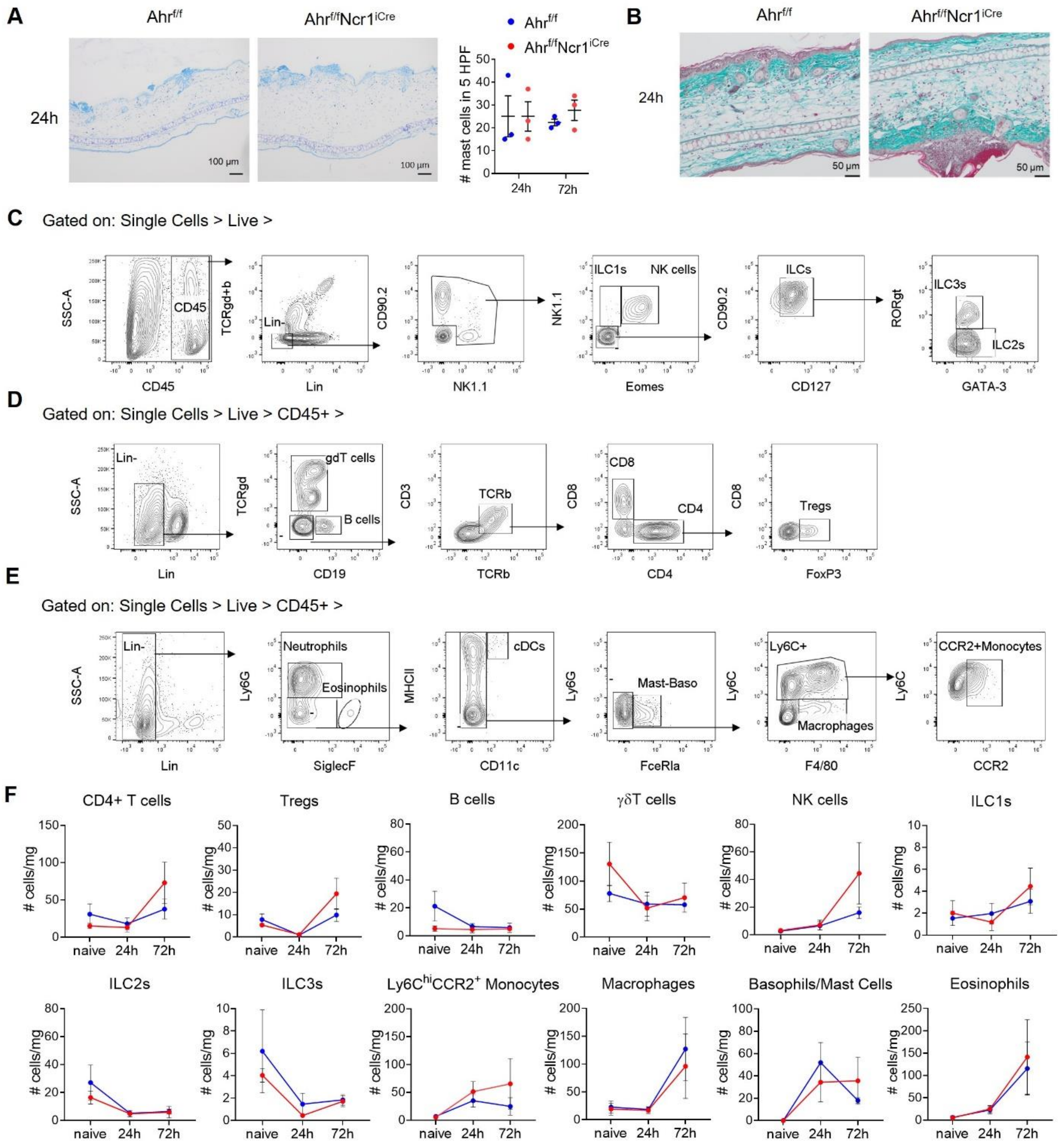


**Figure S1. Characterization of CHS inflammation in $Ahr^{f/f}Ncr1^{iCre}$ mice**

**A)** Representative toluidine-blue staining of the inflamed ears of control $Ahr^{f/f}$ and $Ahr^{f/f}Ncr1^{iCre}$ mice 24h post-challenge (left), and quantification of mast cells in 5 high-power fields (right; n=3 mice per group, mean ± SEM). **B)** Representative trichrome-staining for collagen (aqua-blue) of the inflamed ears of control $Ahr^{f/f}$ and $Ahr^{f/f}Ncr1^{iCre}$ mice 24h post-challenge. Keratin and nuclei are shown in red. **C-E)** Gating strategy applied to identify NK cells and ILCs (C), T cells (D) and myeloid cells (E) in the ear tissue-derived single-cell suspensions by flow cytometry.

Lineage (Lin) in D: Ly6G, SiglecF, FcεRIα, F4/80, Ter119, CD3ε and CD19; in E: Ly6G, SiglecF, FcεRIα, F4/80 and Ter119; in F: CD3ε, CD19, CD127 and NKp46. **F)** Absolute numbers (normalized to tissue weight) of immune populations in the ears of control $Ahr^{f/f}$ (blue) and $Ahr^{f/f}Ncr1^{iCre}$ (red) naive/untreated mice, and 24 and 72h after challenge, analyzed by flow cytometry (n=3-4, mean ± SEM).

*Figure S2*

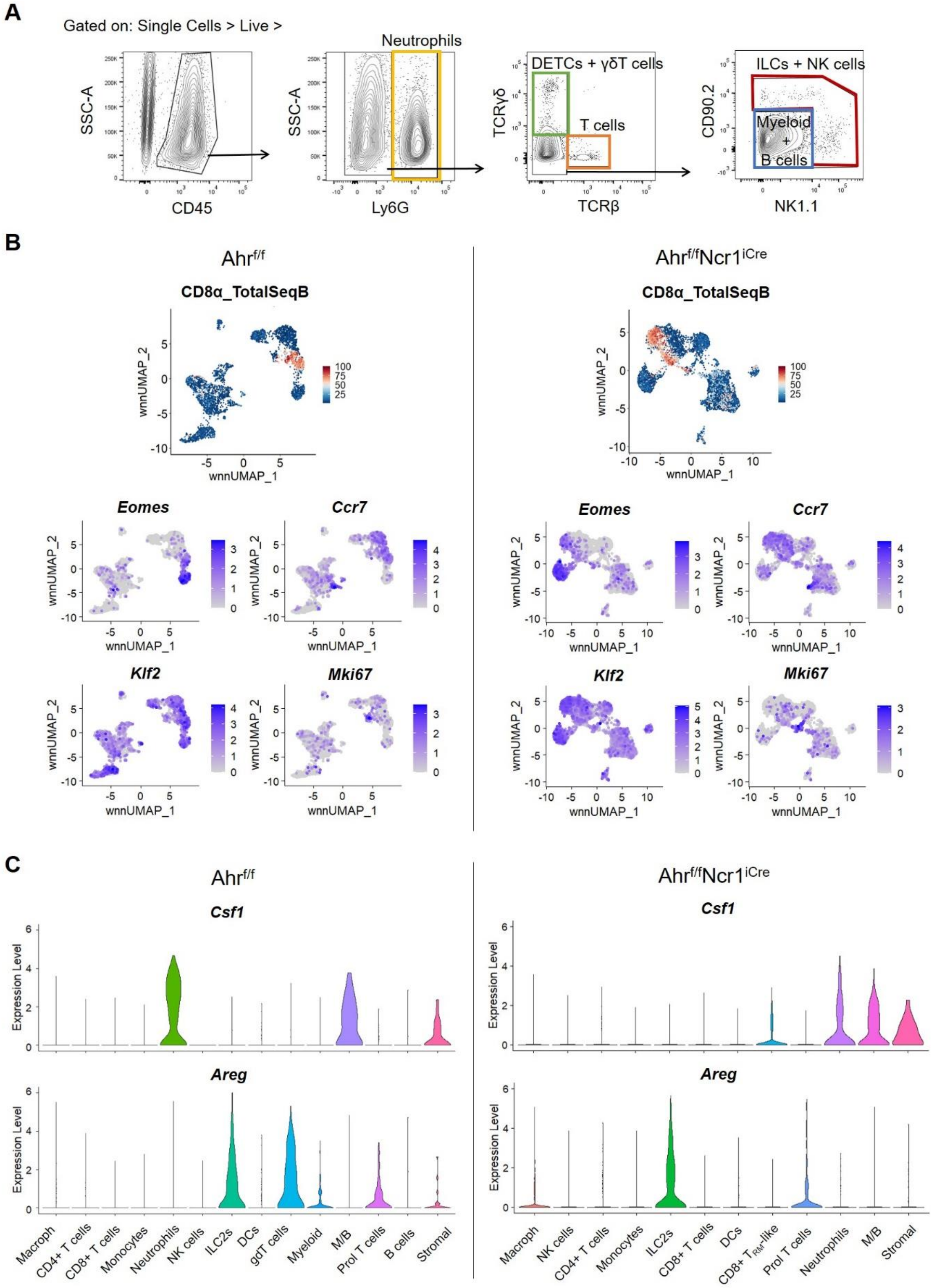


**Figure S2. Cluster-specific expression of selected transcripts.**

**A)** Sorting strategy applied to enrich immune cell populations for multiomic analysis 24h post-challenge. **B)** Detection of cells expressing CD8α surface protein (CD8_TotalSeqB), or selected transcripts (purple) in the inflamed ears of $Ahr^{f/f}$ and $Ahr^{f/f}Ncr1^{iCre}$ mice. **C)** Violin plots showing *Csf1* and *Areg* expression among the identified clusters in the inflamed ears of $Ahr^{f/f}$ and $Ahr^{f/f}Ncr1^{iCre}$ mice. M/B, mast cells and basophils

***Figure S3***

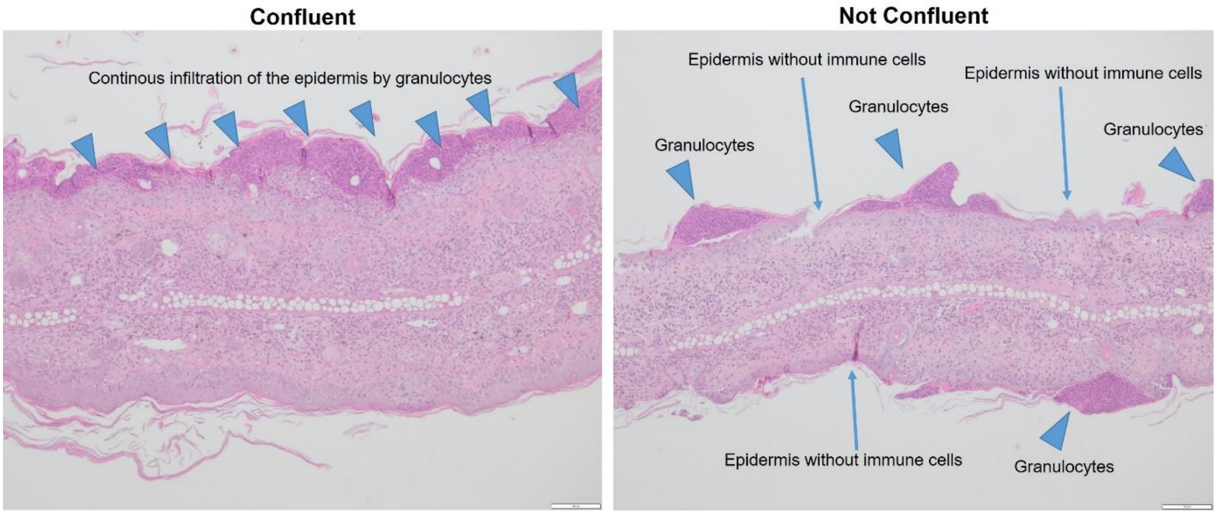


**Figure S3. Confluence of inflammation**

Representative H&E-stained ear sections depicting the presence (left) or absence (right) of confluent inflammatory infiltrate in the challenged ear tissue.
